# Macromolecular crowding is surprisingly unable to deform the structure of a model biomolecular condensate

**DOI:** 10.1101/2022.12.12.520052

**Authors:** Julian C. Shillcock, David B. Thomas, John H. Ipsen, Andrew D. Brown

## Abstract

The crowded interior of a living cell makes experiments on simpler *in vitro* systems attractive. Although these reveal interesting phenomena, their biological relevance can be questionable. A topical example is the phase separation of intrinsically-disordered proteins into biomolecular condensates, which is proposed to underlie the membraneless compartmentalisation of many cellular functions. How a cell reliably controls biochemical reactions in compartments open to the compositionally-varying cytoplasm is an important question for understanding cellular homeostasis. Computer simulations are often used to study the phase behaviour of model biomolecular condensates, but the number of relevant parameters explodes as the number of protein components increases. It is unfeasible to exhaustively simulate such models for all parameter combinations, although interesting phenomena are almost certainly hidden in the jungle of their high-dimensional parameter space. Here we have studied the phase behaviour of a model biomolecular condensate in the presence of a polymeric crowding agent. We used a novel compute framework to execute dozens of simultaneous simulations spanning the protein/crowder concentration space. We then combined the results into a graphical representation for human interpretation, which provided an efficient way to search the model’s high-dimensional parameter space. We found that steric repulsion from the crowder drives a near-critical system across the phase boundary, but the molecular arrangement within the resulting biomolecular condensate is rather insensitive to the crowder concentration and molecular weight. We propose that a cell may use the local cytoplasmic concentration to assist formation of biomolecular condensates, while relying on the dense phase reliably providing a stable, structured, fluid milieu for cellular biochemistry despite being open to its changing environment.

## 1. Introduction

How the myriad biochemical reactions that support cellular life are spatially organised is a fundamental question in cell biology. It is equally crucial for a cell to regulate reactions despite the crowded nature of the cytoplasm. Estimates of the concentration of proteins in eukaryotic cytoplasm range from 100 – 450 mg/ml, which implies that specific macromolecular interactions compete with many random processes involving other molecules.[1-3] Confining reactions inside discrete compartments is one solution to these problems. The nucleus, endoplasmic reticulum, mitochondria, and many other organelles. are wrapped in a phospholipid membrane that segregates their specific biochemistry. In the last decade, an older idea of cellular compartmentalisation has re-emerged as a paradigm for cytoplasmic organisation.[4-8] Dense protein droplets, composed of distinct mixtures of proteins (and often RNA or DNA) have been found to carry out many cellular functions.[9-12]. The proteins that form them lack a minimum-energy folded state, but instead sample a wide ensemble of similar-energy conformations in dilute solution.[13, 14] This has led to their being labelled *intrinsically-disordered proteins* (IDPs). Unlike the organelles mentioned above, these droplets, which are commonly referred to as biomolecular condensates (BCs), have no bounding phospholipid membrane to prevent molecular mixing with the cytoplasm. It is recognised that BCs do not completely isolate their interior from the cytoplasm. Recent work has shown that some kinases respond to molecular crowding in the cytoplasm by condensing into a functional BC that facilitates a signalling pathway to regulate cell volume.[15] Crowding is also known to influence protein folding.[16] Aberrant phase transitions of IDPs are implicated in neurodegenerative diseases, cancer, prion protein diseases, and, more speculatively in ageing.[17-21] Experiments on BCs have advanced to the point that they are being synthetically designed to modulate biochemical and cellular functions,[22-28] and mined for novel drug targets.[29-31] A better understanding of how BCs are coupled (or not) to their crowded environment would assist in deciphering their cellular roles and constructing synthetic BCs.[32-38]

The complexity of living cells has driven research on model condensates, which typically use unphysiologically-high concentrations of one, or at most a few, species of IDP in buffer, although their relevance to living cells has been strongly questioned.[39, 40] An example is provided by *in vitro* experiments on the protein Fused in Sarcoma (FUS), an RNA-binding protein that is implicated in the neurodegenerative disease ALS.[41-44] Computational modelling has also been used to study the phase behaviour of FUS.[43, 45-50] But models that are sufficiently complex to be biologically relevant suffer from a common limitation — their parameter space is huge because many properties of the constituent molecules might *a priori* be important for their behaviour.[51] A popular conceptual model for IDPs is the so-called stickers and spacers model, in which the molecules are regarded as semi-flexible polymers with multiple attractive domains (stickers) connected by weakly-interacting linkers (spacers).[9, 45, 46, 52, 53] When such models aim for one-monomer-per-residue accuracy, they require specifying ∼ 400 interaction parameters just to describe the pairwise forces between the 20 amino acid types.[46]

Further coarse-graining the stickers and spacers model reduces the force field complexity, but still requires assigning values to multiple parameters, including the IDP molecular weight, concentration, backbone stiffness, location of the multiple binding sites, and the interaction energies between all species (solvent, backbone bead, binding sites). We have previously used Dissipative Particle Dynamics (DPD) simulations to explore the phase behaviour and structure of a model biomolecular condensate formed of a single type of IDP modelled on the FUS low-complexity domain (FUS LC).[50, 54-56] Here we ask the simplest question related to the response of a biomolecular condensate to its environment: how does its phase behaviour and internal structure respond to the crowding effect of other macromolecules?

Naively adding additional molecule types as crowding agents multiples the parameters geometrically, rendering it impossible to explore the high-dimensional parameter space of the model in a naively systematic way. We overcome this barrier here by performing multiple, parallel simulations on a novel compute framework.[57, 58] Our goal is to rapidly locate interesting regions of the parameter space and make experimentally-relevant predictions while minimising the computational cost and experimentation time. The simulated FUS-LC molecules are based on previous work,[50] and a soluble polymeric crowding agent (referred to hereafter as the *crowder*) is added that exerts a steric pressure on the IDP molecules (and itself) but is otherwise inert.

Even after these simplifications, the system has a 6-dimensional parameter space — two molecular weights, two concentrations, the IDP self-attraction, and the crowder/IDP repulsion strength — all of which may affect the phase behaviour in a non-trivial way. For simplicity, we set the IDP and crowder molecular weights equal, and initially fix the crowder/IDP repulsion to a high value (see Table 1). This leaves three parameters to be explored in simulations: the crowder and IDP concentrations and the IDP self-attraction.

**Table 1.** Bead-bead conservative force parameters *a*_*ij*_ (in units of *k*_*B*_*T*/*d*_0_) for all bead types. The table is symmetric.

| $a_{ij}$ | W | E | B | F | P |
| --- | --- | --- | --- | --- | --- |
| W | 25 |  |  |  |  |
| E | 25 | $a_{EE}$ | | | |
| B | 23 | 25 | 25 |  |  |
| F | 25 | $a_{EE}$ | 25 | $a_{EE}$ | |
| P | 25 | 80 | 80 | 80 | 80 |

A final difficulty, which is intrinsic to all computational models with many parameters, is our inability to simultaneously visualise multiple states of the model as the parameters are varied. We show that using parallel hardware to accelerate dozens of simulations and displaying the results in a grid of visually-rich simulation snapshots, we can rapidly locate fruitful regions of parameter space. Our results predict that the spatial arrangement and binding of the IDPs within the model condensate is surprisingly insensitive to the crowdedness of its environment even when steric repulsion from the crowder is required for its formation. Biomolecular condensates therefore provide a stable, structured fluid environment for biochemical reactions because the molecular structure of IDPs decouples their internal state (to a certain degree) from their compositionally-varying surroundings.

## 2. Materials and Methods

### 2.1. Dissipative particle dynamics simulations

The Dissipative Particle Dynamics simulation technique (DPD) was invented to study complex fluids such as polymer mixtures, phospholipid membranes, diblock copolymers, etc.[54, 55, 59-61] The simulation source code we use is available on Github.[62] DPD is a coarse-grained molecular technique in which atoms are aggregated into beads that interact via three effective forces. All beads have mass *m*, and the three non-bonded forces between then are soft, short-ranged (vanish beyond a fixed length-scale *d*_0_), pairwise additive, and conserve linear momentum. A conservative force gives each bead an identity:

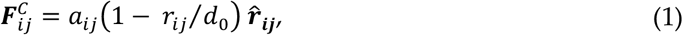

for *r*_*ij*_ < *d*_0_. In this equation, *a*_*ij*_ is the maximum value of the force; ***r***_***ij***_ = ***r***_***i***_ − ***r***_***j***_ is the relative position vector from bead j to bead i, *r*_*ij*_ is its magnitude, and 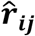is the unit vector directed from bead *j* to bead *i*. Two other non-bonded forces constitute a thermostat that ensures the equilibrium states of the simulation are Boltzmann distributed. The dissipative force is:

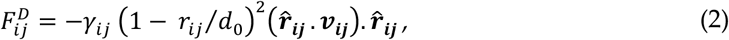

where *γ*_*ij*_ = 4.5 is the strength of the dissipative force, which is the same for all bead types, and ***v***_***ij***_ is the relative velocity between beads *i* and *j*. The random force is:

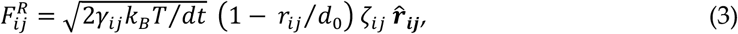

where *k*_*B*_*T* is the system temperature and *ζ*_*ij*_ is a symmetric, uniform, unit random variable that is sampled for each pair of interacting beads and satisfies *ζ*_*ij*_ = *ζ*_*ji*_, ⟨*ζ*_*ij*_(*t*)⟩ = 0, and ⟨*ζ*_*ij*_(*t*)*ζ*_*kl*_(*t*′)⟩ = (*δ*_*ik*_*δ*_*jl*_ + *δ*_*il*_*δ*_*ik*_)*δ*(*t* − *t*′). The factor 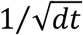in the random force ensures that the discretized form of the Langevin equation is well defined.

Once the required bead types have been specified, they are connected into molecules by tying them together with Hookean springs whose energy function is:

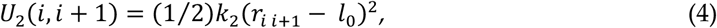

and the spring constant, *k*_2_, and unstretched length, *l*_0_ are here fixed at the values 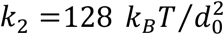and *l*_0_ = 0.5 *d*_0_ for all bond types. Finally, because the peptide chains that form the backbone of the IDPs have a bending stiffness, we add a chain bending potential to the angle *φ* defined by adjacent backbone bead triples (BBB) with the energy function:

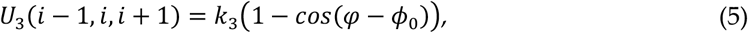

with parameters *k*_3_ = 5 *k*_*B*_*T* and *φ*_0_ = 0.

Three types of molecule are defined in the simulations. The IDPs are semi-flexible, linear polymers containing a set of binding sites (*sticky* bead types E for endcaps and F for internal binding sites) separated by segments of inert backbone beads (B). The crowding agent is a self-avoiding semi-flexible homopolymer composed of a single bead type (P). Solvent molecules are represented by a single bead (W). Both the sticky beads and the backbone beads of the IDP and crowder beads are hydrophilic, and we emphasize that there is no hydrophobic repulsion of the IDPs or crowder from the solvent. In the notation of Ref. 50: 5B6 represents an IDP containing 5 binding sites (including the endcaps) separated by 6 backbone beads (Fig. 1); similarly, 6B10 is an IDP with 6 binding sites separated by 10 backbone beads. The crowder polymers diffuse freely throughout the simulation box exerting an osmotic pressure on the IDPs. Crowder molecules composed of 12, 24 and 48 monomers are indicated by the notation P12, P24, P48 respectively. All non-bonded conservative interaction parameters are given in Table 1. As in previous work,[50] we quantify the attraction of the IDP binding sites by defining a dimensionless parameter *ε* in terms of the conservative force parameter for the binding sites’ self-interaction and interaction with the solvent, namely: *ε* = (*a*_*EW*_ − *a*_*EE*_)/*a*_*EW*_. A value of *ε* = 0 means there is no net attraction between the sticky sites as they have the same interaction with each other as with the solvent beads. Higher values of *ε* indicate stronger attraction. We point out here that bead types E and F have identical interactions in the simulations, but are labelled differently for visual clarity in exploring the phase behaviour.

**Figure 1.**
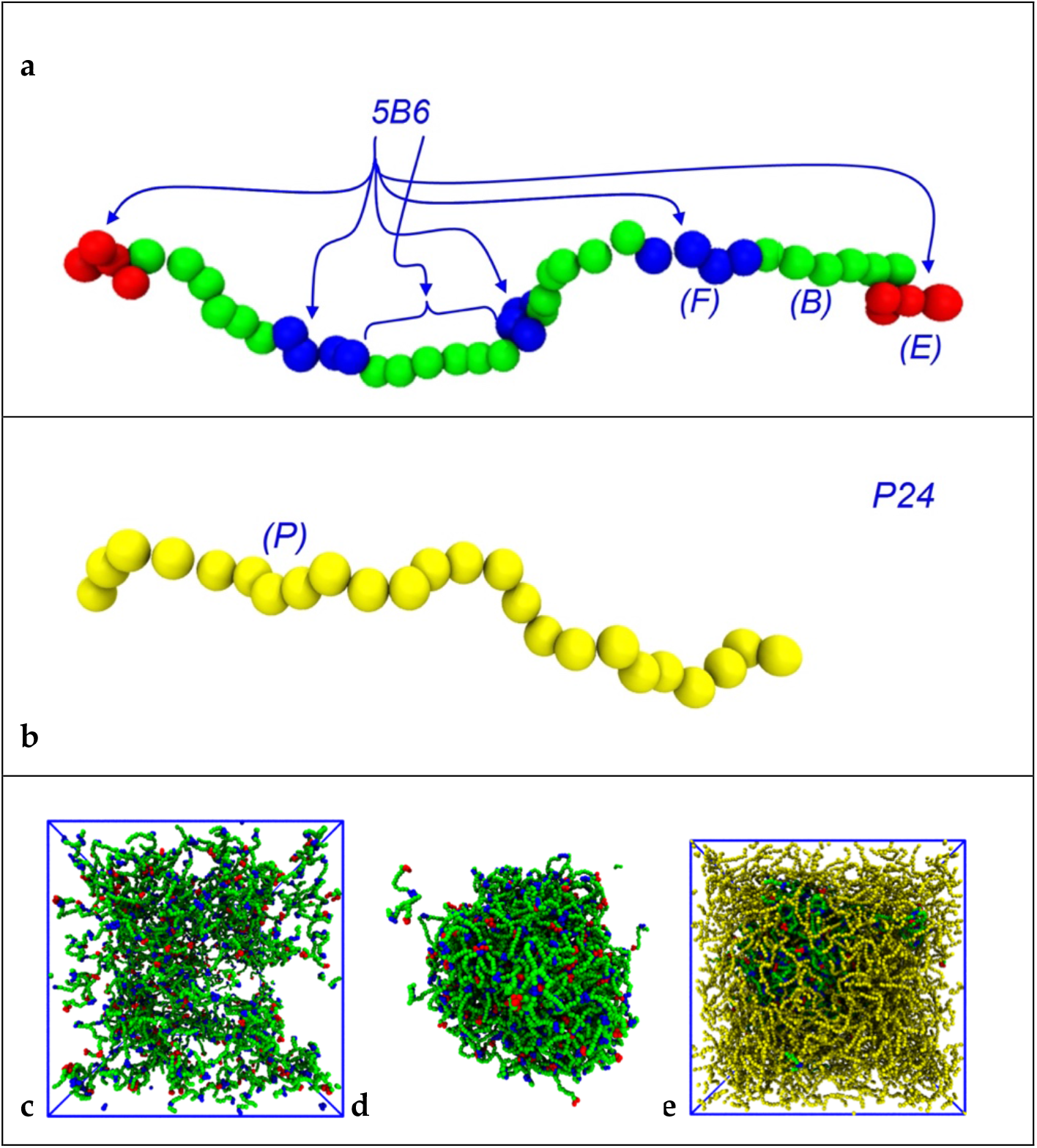
Molecular structure of **a)** 5B6 IDP, and **b)** P24 crowder molecule. The notation 5B6 indicates that there are five binding sites, and adjacent ones are separated by six backbone beads. The endcaps and internal binding sites are coloured differently for clarity only. The crowder polymer beads are yellow and strongly repulsive towards themselves and all IDP bead types (see Table 1). All the IDP and crowder bead types are hydrophilic. **c)** Snapshot of a dispersed system containing 250 IDPs of type 6B10 with no crowder, and **d)** 234 IDPs of the same type in the presence of 468 P48 crowder molecules. **e)** Similar IDP/crowder system with crowder polymers visible. Note the simulation box is not shown and crowder molecules are invisible in panel **d**. Solvent particles are invisible in all figures for clarity.

Simulations take place in a cubical box with dimensions 48 × 48 × 48 (d0)^3^ unless otherwise stated, and periodic boundary conditions are applied. The phase behaviour of the model IDP/crowder system is studied by distributing a given number of IDPs and crowders randomly throughout the simulation box, and filling the remaining space with solvent particles to an average density of density 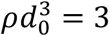 The mass of the beads is set to unity, and the reduced system temperature is *k*_*B*_*T* = 1. Each simulation is run for one million time-steps using an integration step size of 0.02 τ, where 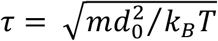is the DPD time-scale. The first half of all simulations is discarded and equilibrium averages are constructed from the second half. Some simulations are extended to two million steps or longer to ensure they reach equilibrium.

### 2.2. POETS

Exploring multiple parameter models is an iterative process involving a human-computer interaction, with the computer performing simulations on a chosen set of parameter values, followed by the human evaluating the results and choosing new sets of parameter values. This type of human-driven parameter space exploration presents a computational and experimental design problems:

1. How do we compute multiple simulations fast enough to allow a semi-interactive parameter-space exploration?
2. What visualisation and workflow support can be created to support a human who wishes to perform such an exploration?

We solved the first problem using multiple high-performance shared-memory multi-core DPD simulation algorithms running in a batch processing system, allowing rapid simulation of multiple points in parallel. The second problem is solved using an automated execution and visualisation workflow wrapped around the parallel simulators, minimising the human effort needed to manage and pre-process results during the search. This approach allowed us to simulate and visualise the results of a 4 × 5 grid of parameters (a 2D parameter space slice) in two hours, while reducing the analysis time needed by the human to five minutes of visual inspection. This should be compared to 7 days if the 20 simulations are executed serially and independently.

Both the simulator and visualisation framework were inspired by a collaborative research project called POETS[58], which is an ongoing project exploring a new computing paradigm called event-triggered computing. In event-triggered computing, the applications and hardware are designed around the frequent exhange of events (small messages) between small asynchronous state machines, rather than the infrequent exchange of large messages between threads (as seen in MPI).

The main idea of POETS is to execute event-triggered applications in custom hardware, allowing applications to scale to thousands of lightweight threads. This idea has previously been applied successfully to DPD simulation, where it was used to tackle simulations of large spatial size.[51] However, while performing that research we uncovered an opportunity and a challenge: the opportunity was that the event-triggered algorithms were also a surprisingly good fit for modern multi-core CPU architectures. The challenge was that there is only one large POETS hardware system, for which there is significant competition for access time.

The fit between event-triggered algorithms and multi-core CPUs resulted in a very fast multi-core shared memory DPD simulator. The technical details are not the focus of this paper, but the main ideas are that it uses very fine-grain spatial domain decomposition so that each thread manages a unit-volume cell, with movement/copying of beads between domains rather than building and maintaining edge lists. The event-driven nature of the algorithm makes it particularly amenable to SIMD vectorisation, reduces cache traffic, and allows for efficient shared memory multi-threading.

We do not make claims about the relative efficiency of this approach for all computational problems, but for the experiments performed in this work using the Iridis data centre (https://www.southampton.ac.uk/isolutions/staff/iridis.page) this resulted in a 2x speed-up compared with the industry-standard code LAMMPS (https://www.lammps.org) in a single 64-core system. Two important aspects of this setup were:

- It allowed experiments to complete in under two hours, which allows them to be submitted to the “fast” low-latency queue; and
- It means we do not need to use GPUs to achieve low-latency, which is useful because GPUs are less common in many HPC clusters, and heavily subscribed by chemists, physicists, and machine learning researchers. In our experience it typically takes more than 2 hours for a GPU job to even start running, even though it may execute faster once scheduled.

So the result of having a very fast multi-core shared-memory is that multiple jobs can be issued and completed very quickly. Clearly these exact figures are not true of all HPC systems, but it does address two common problems: GPUs are a very contested resource, and jobs need to have short run-times in order to be scheduled with low latency.

This multi-core oriented approach allows 20 independent 48×48×48×1M simulations to complete in under 2 hours, which is close to interactive – in principle a researcher could perform four simulate-observe-decide iterations in a working day. In practise we found that there was substantial overhead due to preparing simulation inputs, collecting outputs, and then post-processing them into a useable form that a human could analyse. Simple tasks like ordering and organising images take large amounts of time, involving opening multiple files and trying to arrange them in some way to support interpretation and analysis. Creating and submitting the next batch of simulations is then another time consuming steps.

Through experimentation we found that the most effective approach – at least for our current purposes - is to explore a plane of two parameters, and then automatically arrange the simulation outputs into a visual 2D grid (e.g., see Figure 2). In most cases this allows the human to immediately explore and interpret the results. If the current parameter ranges are not exposing any behaviour of interest, a new set of parameters can be found and re-submitted for exploration. If an interesting set of parameters has been hit, then either more detailed simulations can be performed in the same area, or the human can take the simulation results and start to explore them using other approaches for statistical and visual analysis.

**Figure 2.**
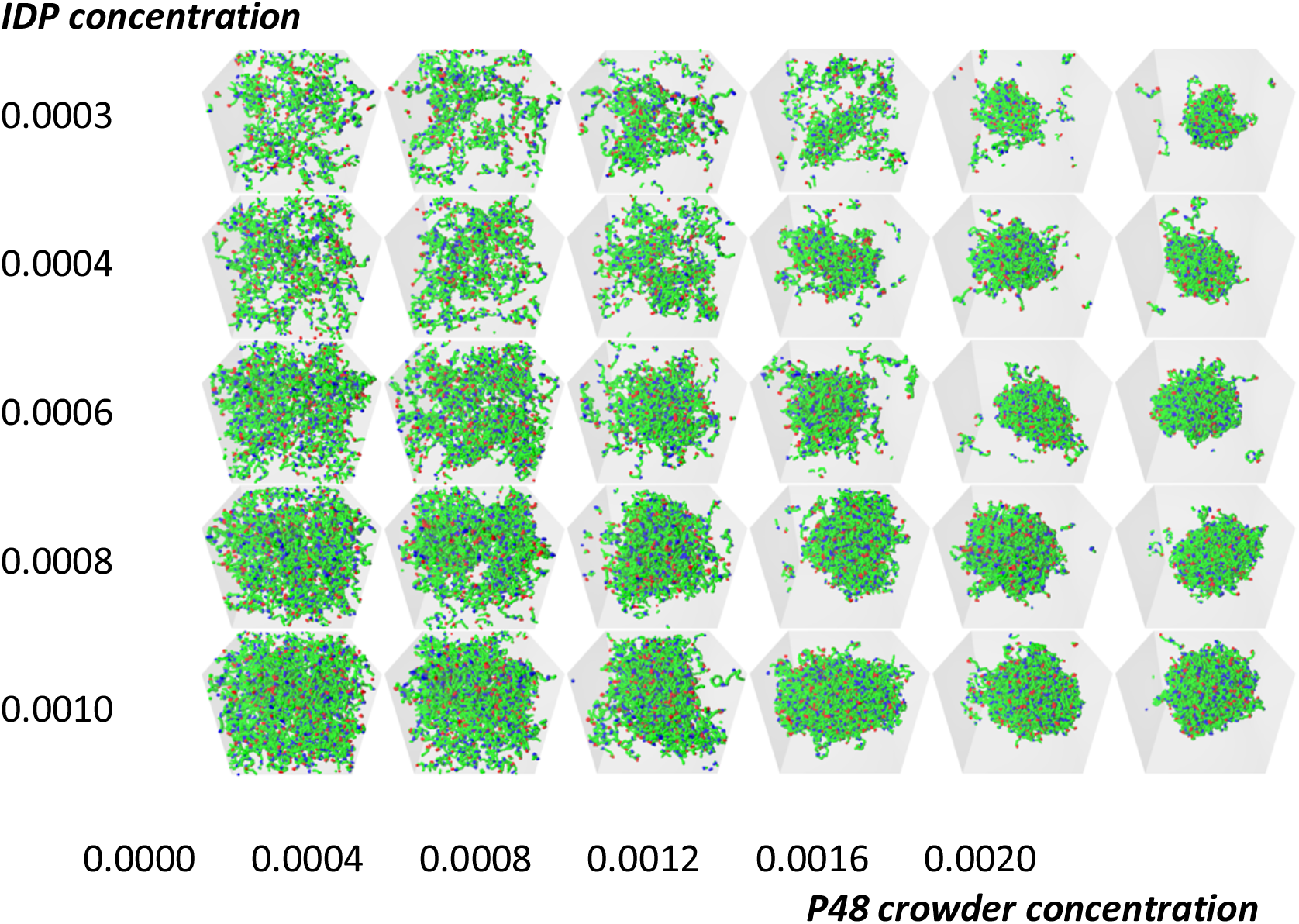
Illustrative grid of snapshots of 6B10 molecules with constant affinity *ε* = 0.6 in the [Crowder]-[IDP] plane. The rows have constant IDP concentrations (top to bottom): 0.0003, 0.0004, 0.0006, 0.0008, 0.001, and the P48 crowder concentration increases across the columns (left to right): 0.0, 0.0004, 0.0008, 0.0012, 0.0016, 0.002. Increasing the crowder concentration (top row) drives the system across its phase boundary producing a dense droplet surrounded by a dilute phase. Examining the lower rows of the grid reveals that the crowder concentration required to drive phase separation decreases with increasing IDP concentration (bottom row). Solvent and crowder molecules are invisible in all snapshots for clarity.

The overall workflow we developed for the results used in this paper is:

1. *Manual*: Define a model with multiple parameter dimensions to sweep
2. *Manual*: Construct a parameterised scenario generator which can instantiate the model for specific parameter values
3. Perform human-computer collaborative search:
  a. *Manual*: identify two interesting parameter dimensions and ranges; pick X points for one parameter and Y for the other and generate the XxY concrete scenarios to simulate
  b. *Automatic*: Simulate scenarios in parallel on multiple machines in a HPC system
  c. *Automatic*: Collect outputs and produced tiled XxY images and videos
  d. *Manual*: inspect tiled images to explore parameter response. If necessary, go back to step 2.a
4. *Manual*: Explore and analyse the results in more detail

The two manual bottleneck tasks in this iterative process are 3.a and 3.d : picking the parameter values at the start of each iteration, and manually investigating the results. Using a set of scripts we reduced 3.a down to the point where the user submits a zip file of simulation scenarios. Each simulation scenario in the zip is an Osprey DPD simulation description file, with a file-name prefix indicating its position within an integer grid. At step 3.d the user then receives back a zip file containing periodic snapshots of each simulation’s state, while also receiving automatically assembled grids of rendered images showing the spatial variations (see Figure 3). In many cases it is sufficient to simply open the assembled grid for the image slice to get a sense of the parameter dependency, and immediately produce the next set of scenarios.

**Figure 3.**
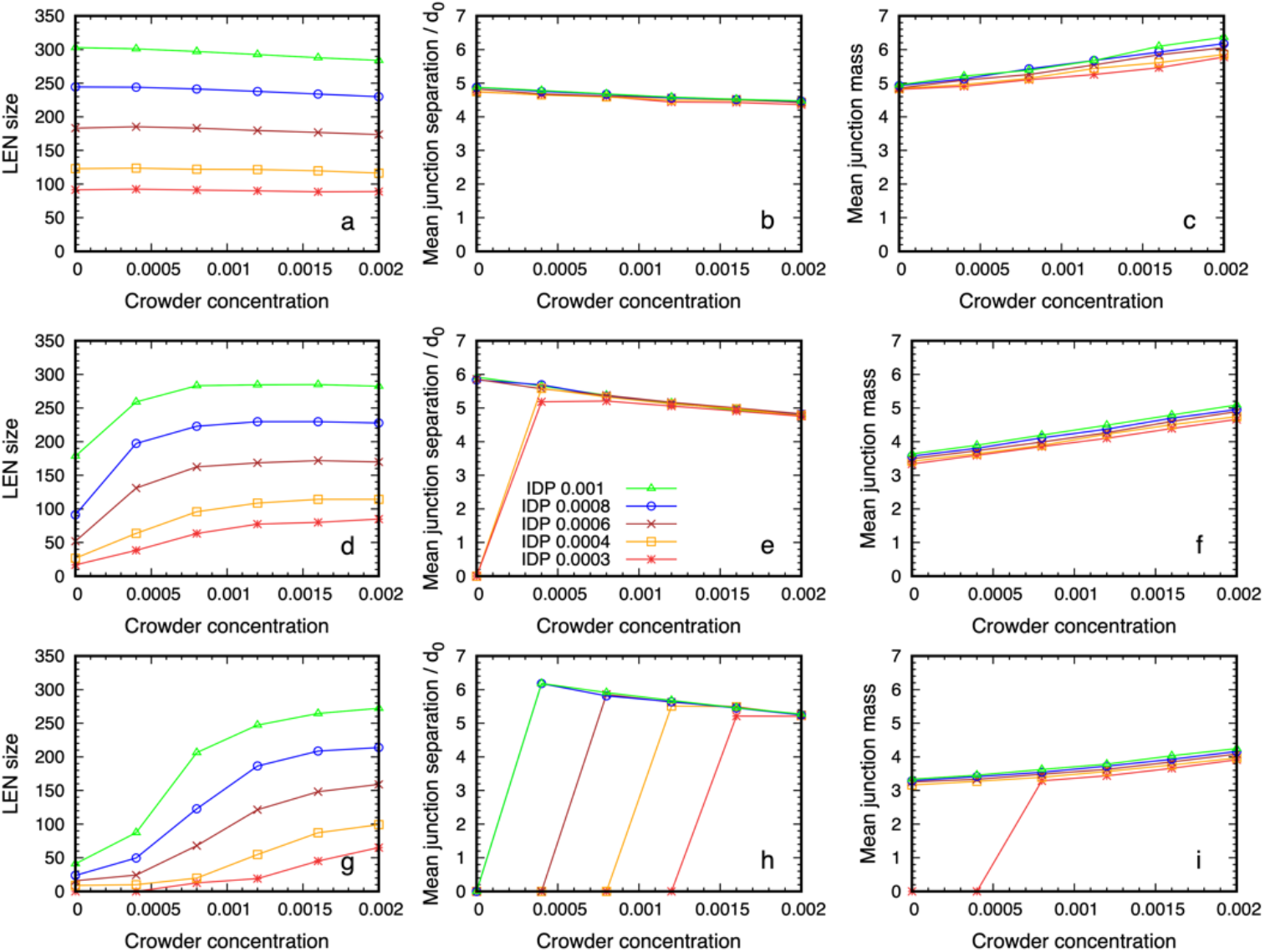
Structural data on the dense phase of 6B10 IDPs with affinities *ε* = 0.6, 0.68, 0.76 (bottom to top) as a function of the P48 crowder concentration for several values of IDP concentration. Panels a, d, g in the first column show that the number of IDPs in the Largest Equilibrium Network (LEN) increases with IDP concentration, and that lower affinity requires a higher crowder concentration before the dense phase appears. The middle column (b, e, h) shows that the separation between the junctions at which IDP binding sites meet is independent of their concentration, and decreases slowly with increasing crowder concentration. The final column (c, f, i) shows that the mean number of IDPs binding at the junctions is independent of the IDP concentration but increases weakly with increasing crowder concentration. Note that if the LEN does not exist, data points lie on the abscissa.

The automated tools supporting steps 3.b and 3.c are implemented using a combination of the POETS DPD software simulator, python scripts to assemble image grids, and a set of shell-scripts to manage them. The grids are executed on the Iridis-5 HPC cluster at the University of Southampton, with one 64-core AMD node per simulation. Each node is able to complete a 48×48×48×1M simulation in less than 2 hours, requiring about 3.3×10^11^ bead-steps. A grid of 4×5 simulations typically completes 6.6×10^12^ bead-steps in 2 hours.

This work-flow could usefully be replicated using other tools – for example, it can be performed using Osprey DPD, though it would slow the iteration time down to at most one iteration per day. It could also be implemented using a GPU accelerated simulator such as LAMMPs, and in an HPC system with abundant GPUs this could be a good solution. However, GPU nodes are still often a small subset of total compute nodes in many HPC pools, so even if simulation tasks execute more quickly, the tasks may spend a long time waiting before execution.

To give a concrete example: in the Iridis-5 system there are 572 multi-core CPU compute nodes, but only 20 GPU nodes – a not uncommon balance for a large institution-wide HPC pool for batch computing. As a result there is intense competition for GPU access time due to physicists, chemists, engineers, and machine learning researchers all wanting to run long multi-hour tasks, so GPU tasks may spend 10-20 hours waiting in the submission queue. To complete a grid of 20 GPU tasks might take 2-3 days, even though it only takes 20 hours of on-node time, eliminating the interactive aspect of the parameter space exploration. By using fast multi-core software simulators we can get all tasks executing rapidly, exploiting the large number of CPU compute nodes available for short-lived tasks, and complete all 20 tasks in a semi-interactive two hour timescale.

## 3. Results

### 3.1. Crowding assists phase separation of IDPs with sub-critical affinity

Phase separation of IDPs is driven by transient attractive interactions between specific amino acids, such as arginine and, tyrosine, and non-specific electrostatic and hydrophobic interactions between residues.[63] It is assisted by the relatively small entropic cost when proteins move from a dilute phase into the dense phase, as demonstrated by FUS that retains high conformational flexibility inside its dense phase.[42, 43] In previous work, we showed that model IDPs of the type shown in Figure 1 spontaneously phase separate when the affinity of their binding sites exceeds a theshold that depends on the number and location of the binding sites.[50] When their affinity is reduced below the critical value, no spontaneous phase separation occurs. Here, we first explored how the presence of a crowding agent influences the phase separation of a model IDP whose affinity is below the critical value.

The model IDP has 6 binding sites separated by 10 backbone beads, and is represented by the notation 6B10. Previous work shows that the critical affinity for such molecules lies between *ε* = 0.68 − 0.74.[50] We set the binding site affinity to *ε* = 0.6, which is below the critical value, so the IDPs are unable to phase separate spontaneously. The crowding polymer was P48 and all other parameters were fixed as described in the Methods section. This left a two-dimensional parameter space, the IDP and crowder concentrations, which can conveniently be displayed as a two-dimensional array of results. Figure 2 shows a grid of snapshots taken from 30 simultaneous simulations of 10^6^ time steps performed using POETS-DPD (see Methods). The rows have constant IDP concentration increasing towards the bottom, and the columns have constant crowder concentrations, increasing to the right. The crowder molecules are invisible in the figure for clarity, but Supplementary Figure S1 shows the complete systems. The top row shows the general trend in the observed equilibrium states as the crowder concentration is increased. In the absence of crowder, the IDPs are dispersed (top left image), while increasing the crowder concentration eventually drives the system across the phase boundary, resulting in a phase separated droplet surrounded by a dilute phase (top right image). Each lower row shows a higher IDP concentration, and the same general trend is seen. However, when the IDP concentration is sufficiently high, they already form a loose network that spans the box in the absence of the crowder, a result previously reported in the literature.[45, 50, 56] Visually comparing all the snapshots in the grid shows that higher concentrations of IDP phase separated at lower crowder concentrations. When the same IDP/crowder concentrations were simulated with higher affinities *ε* = 0.68, 0.76, the boundary between the dispersed and phase separated states shifted to lower crowder concentrations with increasing affinity (see Fig. 4 and Supplementary Figure S2).

**Figure 4.**
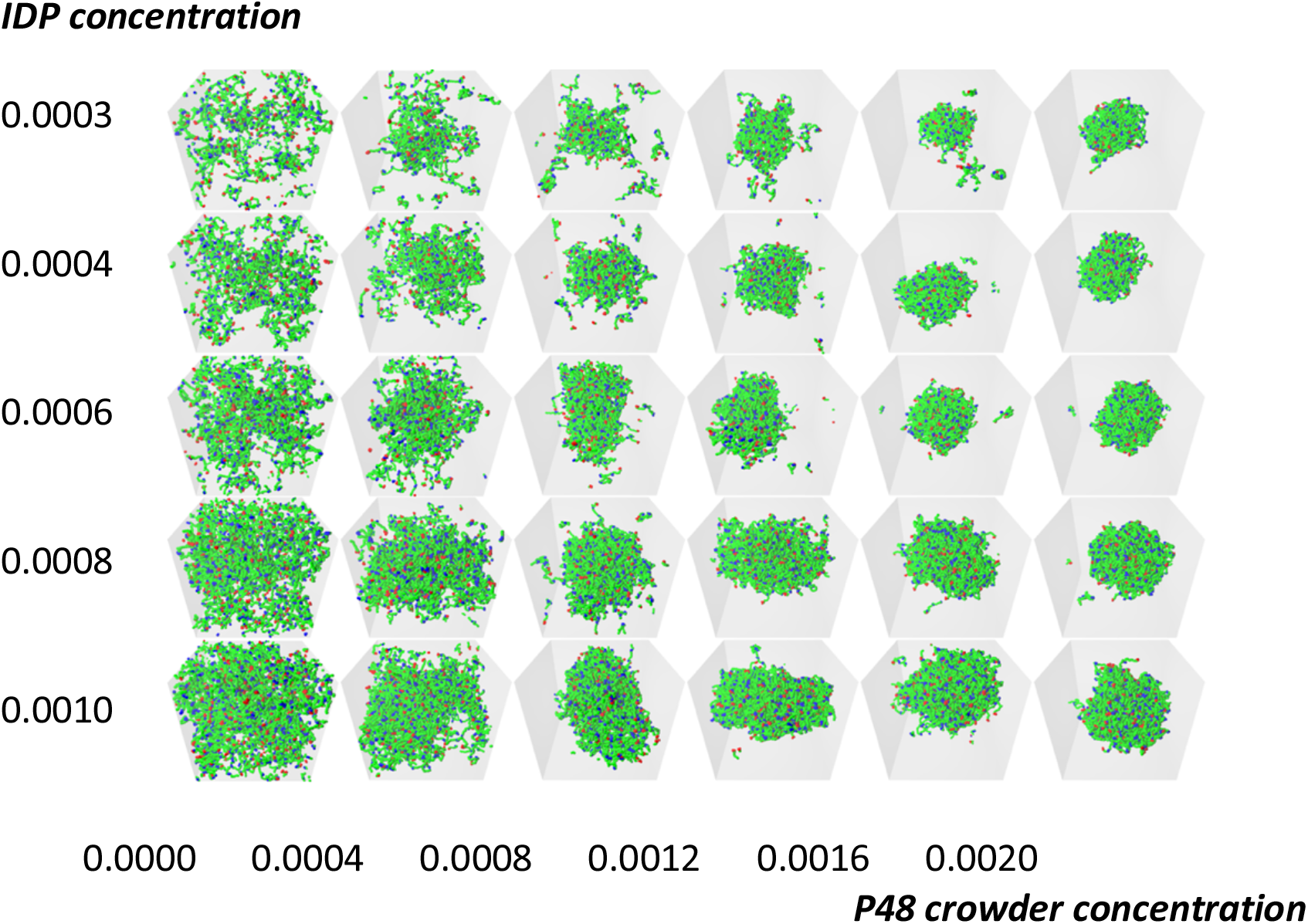
Illustrative grid of snapshots of 6B10 molecules with affinity *ε* = 0.68 in the [Crowder]-[IDP] plane for P48 crowder polymers. As in Fig. 2, the rows have constant IDP concentrations (top to bottom): 0.0003, 0.0004, 0.0006, 0.0008, 0.001, and the crowder concentration increases across the columns (left to right): 0.0, 0.0004, 0.0008, 0.0012, 0.0016, 0.002. The binding site attraction is stronger than in Fig. 2, but still too weak to drive phase separation in the absence of crowder as seen in the first column.

It might be expected that a sufficient concentration of the repulsive crowder could drive phase separation of IDPs whose affinity is low or even zero. This would represent a generic polymer phase separation driven by the steric repulsion between the molecules, and described by Flory-Huggins theory.[64] In particular, it would not depend on the self-affinity and number of IDP binding sites, and therefore the heterogeneous spatial organisation of the molecules in the dense phase observed in earlier work might be eliminated.[50] To test this possibility we ran control simulations in which the binding affinity of the IDPs was initially set to a value known to drive phase separation (*ε* = 0.68), and subsequently removed after 500,000 time steps (*ε* = 0). Supplementary movie SM1 shows the evolution of the system with [IDP] = 0.0006 (180 molecules) and [Crowder] = 0.0012 (361 molecules), which corresponds to grid element 3,4 in Fig. 2. When the affinity was removed, the dense phase dissolved, showing that the presence of the crowder alone was unable to drive phase separation for this combination of IDP/crowder molecular structure and concentrations. This does not preclude that a higher crowder concentration could drive phase separation. Supplementary movie SM2 shows the result for grid element 3,6 in Fig. 2 showing that the higher crowder concentration of 0.002 (583 molecules) was sufficient to keep the IDPs from dispersing in the absence of any self-attraction between the IDPs. However, when we analysed this dense phase, the binding sites of the IDPs did not form junctions, and the IDPs are not in a connected network.

We calculated the crowder concentrations as follows. The densest case in the grid, which corresponds to the rightmost column in Figure 3, has between 568 and 594 molecules of the P48 crowder in the simulation box (the precise value depends on the number of IDP molecules). The molecular weight of the crowder was chosen to be equal to that of the IDP, which represents FUS-LC, and its concentration was calculated using the method of Ref. 50. This gave a concentration of 7 mM, equivalent to 120 mg/ml, which is in the range of estimates for the eukaryotic cytoplasm of 100 – 450 mg/ml.[1, 2]

### 3.2. Quantitative properties of condensate structure are insensitive to crowder concentration

We next explored whether the internal organization of the IDPs in the dense phase is modified when the phase separation is assisted by the crowding agent compared to when it forms spontaneously (i.e., when the IDPs have a higher affinity). We found previously that the dense phase has a heterogeneous structure in which the binding sites of the IDPs meet at spatially-discrete junctions whose separation depends on the spacing of the binding sites but is less sensitive to their affinity (cp Fig. 3 in Ref. 50). Further, the average number of IDPs that span two junctions increases with increasing binding site affinity and decreasing separation. These junctions are transient because the IDPs continually fluctuate and diffuse through the dense phase.

We show in Figure 4 the quantitative properties of the dense phase of 6B10 IDPs for three values of the affinity, *ε* = 0.6, 0.68, 0.76, near the critical value as a function of the crowder concentration. All the systems in the top row (panels a, b, c) are phase separated, even in the absence of the crowder, because of their high self-affinity (the grid of snapshots for this affinity, *ε* = 0.76, are shown in Supplementary Figure S2). Panels g, h, i along the bottom row correspond to the array of simulations in Figure 3, and panels d, e, f in the middle row correspond to those in Fig. 4. The first column shows the number of IDPs that are connected together to form the Largest Equilibrium Network (LEN). This is defined at each sampling point of the simulations as the largest set of IDP molecules that are simultaneously connected by their binding sites. The continual exchange of molecules between the dilute and dense phases causes the LEN to fluctuate over time, and we aveage over many samples to obtain its equilibrium size and properties. We recalculated the LEN for each sample used in the quantitative analysis to minimise the influence of small clusters and surface effects as described previously.[50] Whereas the size of the LEN for high affinities (4. a) is independent of the crowder concentration, lower affinities (4. d, g) required a higher crowder concentration before the LEN became stable. Note that values of the observables near zero means there were too few IDPs in the LEN to calculate equilibrium averages, i.e., almost all the IDP molecules are dispersed. The fraction of IDPs in the dense phase increased with crowder concentration, until it contained all the IDPs at the highest concentration.

The second column shows the mean junction separation within the condensed phase as a function of the crowder concentration for all IDP concentrations studied. The separation is clearly independent of the IDP concentration (and therefore the droplet size) and the crowder concentration over the range where the dense phase is stable. Comparing the junction separation in panels 4. b, e, and h shows that it is insensitive to the binding site affinity and crowder concentration over the studied range apart from a slow decrease which is less than the size of the monomers forming the IDP. Finally, the mean junction mass is insensitive to the IDP concentration but shows a small systematic increase with increasing crowder concentration (Fig. 3 c, f, i). Taken together, these results show that IDPs with weaker affinities require higher crowder concentrations to phase separate. The mean junction separation in the dense phase is largely insensitive to the IDP and crowder concentrations, while the number of IDPs binding at the existing junctions increases slowly with increasing crowder concentration.

### 3.3. Dense phase structure is partially decoupled from the crowder molecular weight and enthalpic repulsion from the IDPs

#### 3.3.1. Reducing the crowder volume fraction shifts crowding-enhanced phase separation but does not greatly perturb the dense phase structure

The previous section showed that the structural organisation of the IDPs in the equilibrium dense phase is independent of the IDP concentration (once the dense phase is stable), and only weakly dependent on the crowder concentration. We next probed the dependence of this structure on the molecular weight of the crowder. Because we expected the reduced volume fraction of shorter crowders to impose a smaller osmotic pressure on the IDPs, we increased the IDP affinity from *ε* = 0.6 to *ε* = 0.68. The baseline results using P48 crowders are shown in Figure 4. The first column shows that the higher self-affinity is still too weak to drive phase separation in the absence of crowder, but the IDPs phase separate on increasing crowder concentration as observed in Fig. 2.

We then repeated these simulations with the size of the crowding polymers reduced from 48 to 24 monomers, keeping their number fractions for each grid element the same as before (Figure 5). This effectively reduces their volume fraction by a factor of two. (Supplementary Figure S3 shows the grid with the P24 crowder molecules visible). Comparing Figures 4 and 5 shows that the phase transition of the IDPs is shifted to a higher crowder concentration in the presence of P24 crowders as P48, because more IDPs move from the dense to the dilute phase, but the droplet is still formed.

**Figure 5.**
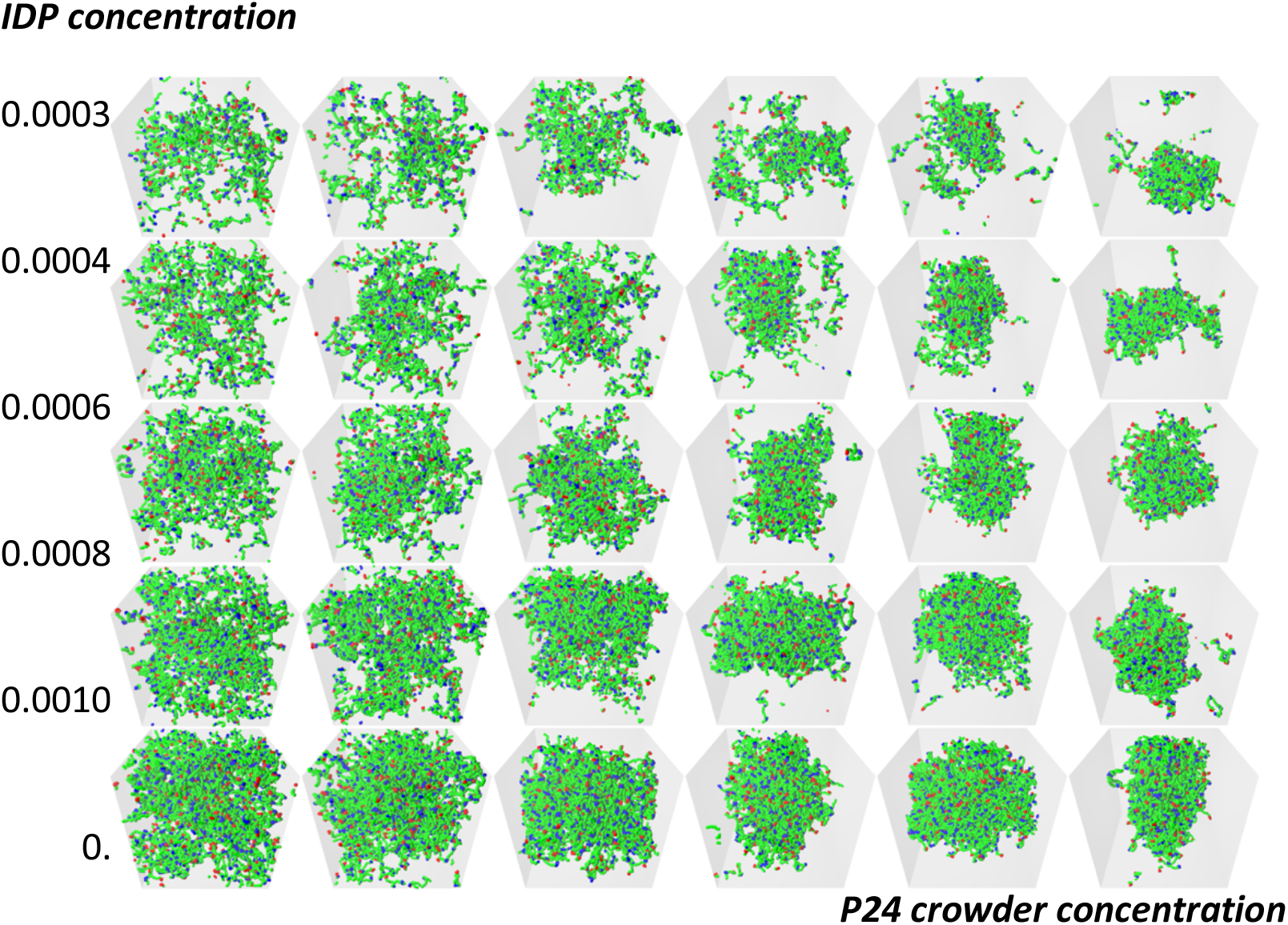
Qualitative effect of reducing the length of the crowder molecules on the dense phase stability of 6B10 molecules with constant affinity *ε* = 0.68. Replacing P48 crowders by P24 with the same number fraction halves the monomer volume fraction of the crowders. The IDPs phase separate at higher crowder concentrations, and more IDPs are in the dilute phase compared to in Figure 4.

We measured the quantitative structure of the dense phase in the presence of P24 crowders, and show the results in Figure 6. For the IDPs with the highest affinity, the dense phases are almost identical for the two types of crowding polymer. When the affinity is lowered, the most noticeable effect is that a higher crowder concentration is needed to drive the phase separation of the IDPs, which is intuitively expected. Once the dense phase appears, its quantitative structure is only weakly dependent on the crowding polymer length. For the lowest affinity case (Panels 6. g, h, i), the mean junction separation increased by approximately 20% and and the mean junction mass decreased by 10-15%. This shows that the spatial organization of the IDPs inside the dense phase is largely decoupled from the molecular weight of the crowders when their volume fraction is constant.

**Figure 6.**
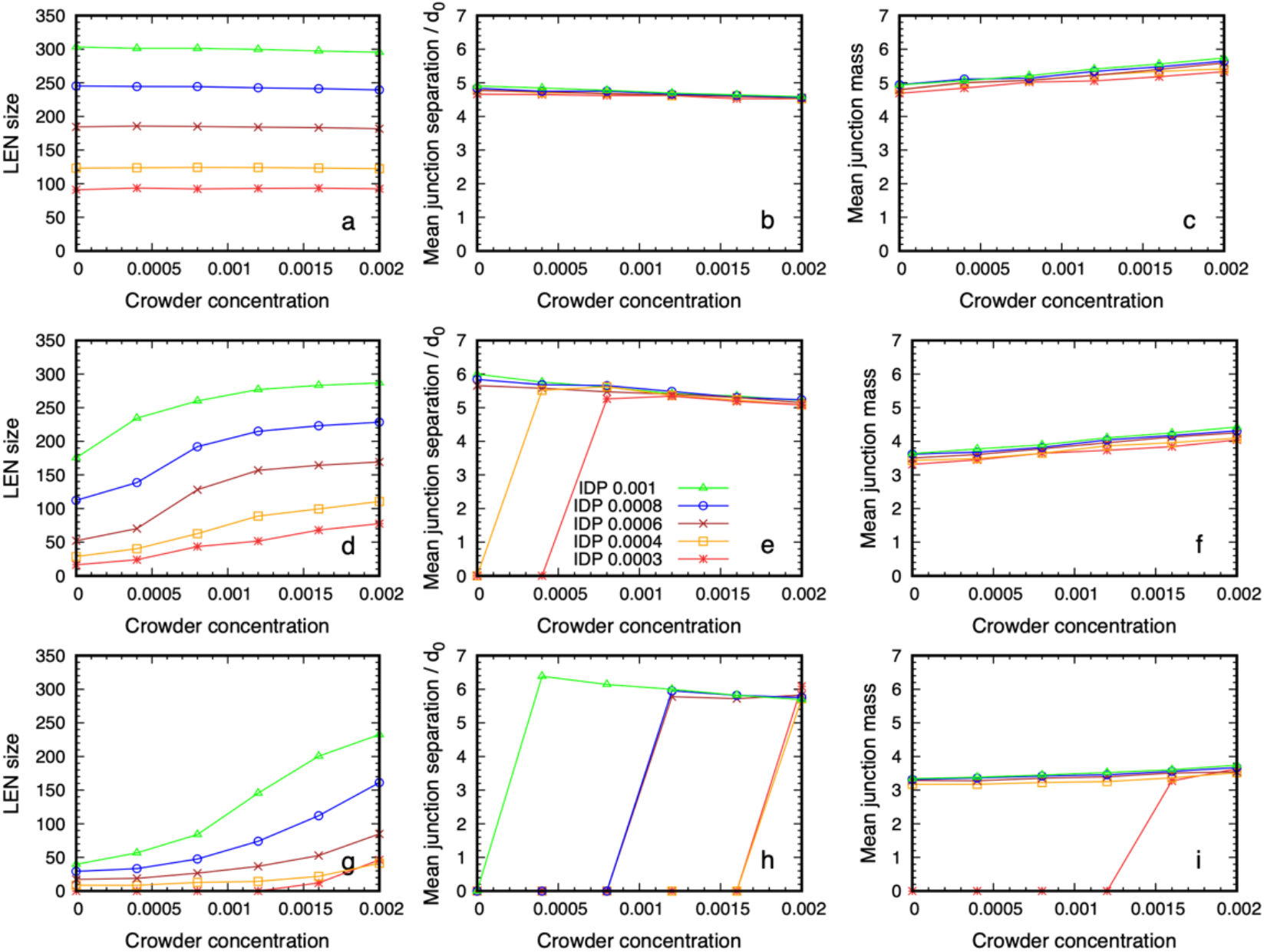
Structural data on the dense phase of 6B10 IDPs with affinities *ε* = 0.6, 0.68, 0.76 (bottom to top) as a function of the P24 crowder concentration for several values of IDP concentration. Panels a, d, g in the first column show that the number of IDPs in the Largest Equilibrium Network (LEN) increases with IDP concentration, and that a lower affinity requires a higher crowder concentration before the dense phase appears. The middle column (b, e, h) shows that the separation between the junctions at which IDP binding sites meet is independent of their concentration, and decreases slowly with increasing crowder concentration. The final column (c, f, i) shows that the mean number of IDPs binding at the junctions is also independent of the IDP concentration but increases weakly with increasing crowder concentration. Note that if the LEN does not exist, data points lie on the abscissa.

We next replaced the P24 crowders by P12 and qualitatively checked whether the dense phase remains stable. We selected the system in the third row and fourth column of Figure 4 as a typical case. The IDP and crowder number fractions were 0.0006 and 0.0012 respectively (corresponding to 180 IDPs and 361 crowders). The conservative repulsive parameter between the crowder monomers and IDP monomers was kept at *a*_*Px*_ = 80, where x stands for all bead types except solvent. (cp. Table 1). We performed separate simulations of the P24 crowders with number fractions 0.0012 and 0.0024, and P12 crowders with number fractions 0.0012 and 0.0048. Figure 7. a and b show that P24 crowders at the same number fraction as P48 reduces the stability of the droplet, and more IDPs migrate to the dilute phase. Keeping the P24 volume fraction equal to that of the P48 crowder restores the droplet’s stability. Supplementary movie SM3 shows the evolution of the P24 system over 10^6^ time steps for the same number fraction and same volume fraction. Figure 7. c and d show the same result for P12 crowders: the droplet dissolves due to the reduced crowding effect of the P12 molecules at the same number fraction, but its stability is restored when their number fraction is increased by a factor of four. Supplementary movie SM4 shows the evolution of the P12 system over 10^6^ time steps for the same number fraction and volume fraction.

**Figure 7.**
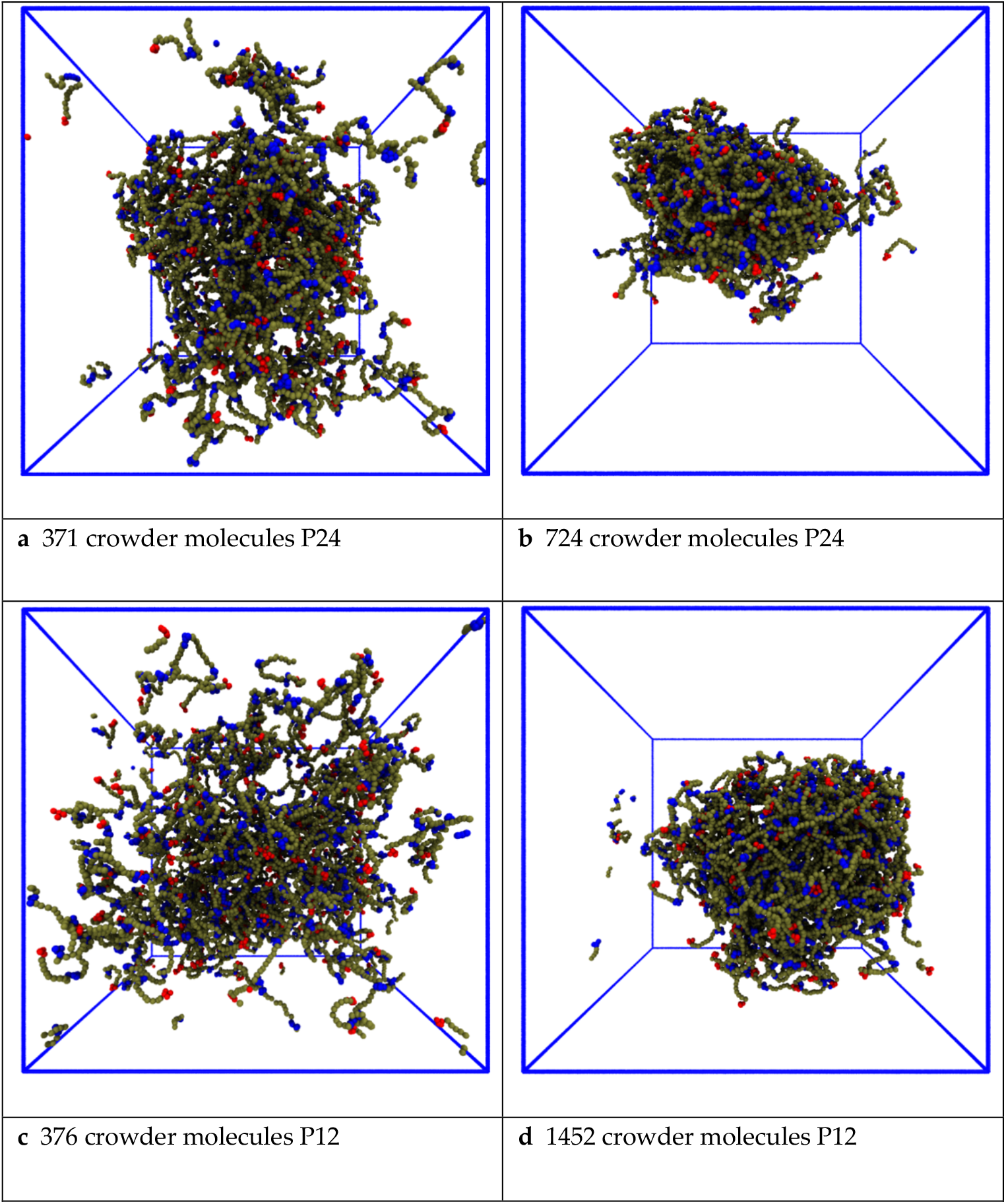
Effects of reducing the length of the crowder molecules on the dense phase stability. **(a)** Replacing P48 crowders by P24 at a constant number fraction halves the volume fraction. The droplet remains phase separated but more IDPs move into the dilute phase. **(b)** The droplet becomes stable again when the number fraction of P24 crowders is increased by a factor of 2, thereby restoring their volume fraction to the original value. **(c)** The droplet dissolves when the P48 crowders are replaced by P12 at constant number fraction. **(d)** The droplet is stable again when their number fraction of P12 crowders is increased by a factor of 4, thereby restoring their volume fraction.

#### 3.3.2. Reducing the enthalpic repulsion of the IDPs from the crowder molecules leaves the dense phase stable

Next, we investigated the impact of reducing the enthalpic repulsion between the IDPs and crowder molecules on the phase separation. We again started with the system in the third row and fourth column of Figure 4. The baseline repulsion of *a*_*Px*_ = 80 between the crowder monomers and all other bead types x was reduced to *a*_*Px*_ = 50, 35 25 at 500,000 steps in independent simulations. Figure 8 shows snapshots of the systems after 2 10^6^ time steps to ensure that the IDPs have time to disperse. The IDPs do not phase separate for this combination of concentration and affinity in the absence of the crowder (panel 8a) nor when the repulsion between the crowder and IDP polymers is the same as the IDP self-repulsion (panel 8b). The crowding agent was able to assist the phase separation when its repulsion from the IDPs was greater than *a*_*Px*_ = 35, a value not far above the IDP’s self-repulsion (panels 8c and d). This shows that the phase separation is not greatly sensitive to the enthalpic repulsion between the IDPs and the crowding agents as long as it is above a minimal threshold (Supplementary movie SM5 shows the evolution of the systems shown in panels 8 b and d for the second 10^6^ time steps.)

**Figure 8.**
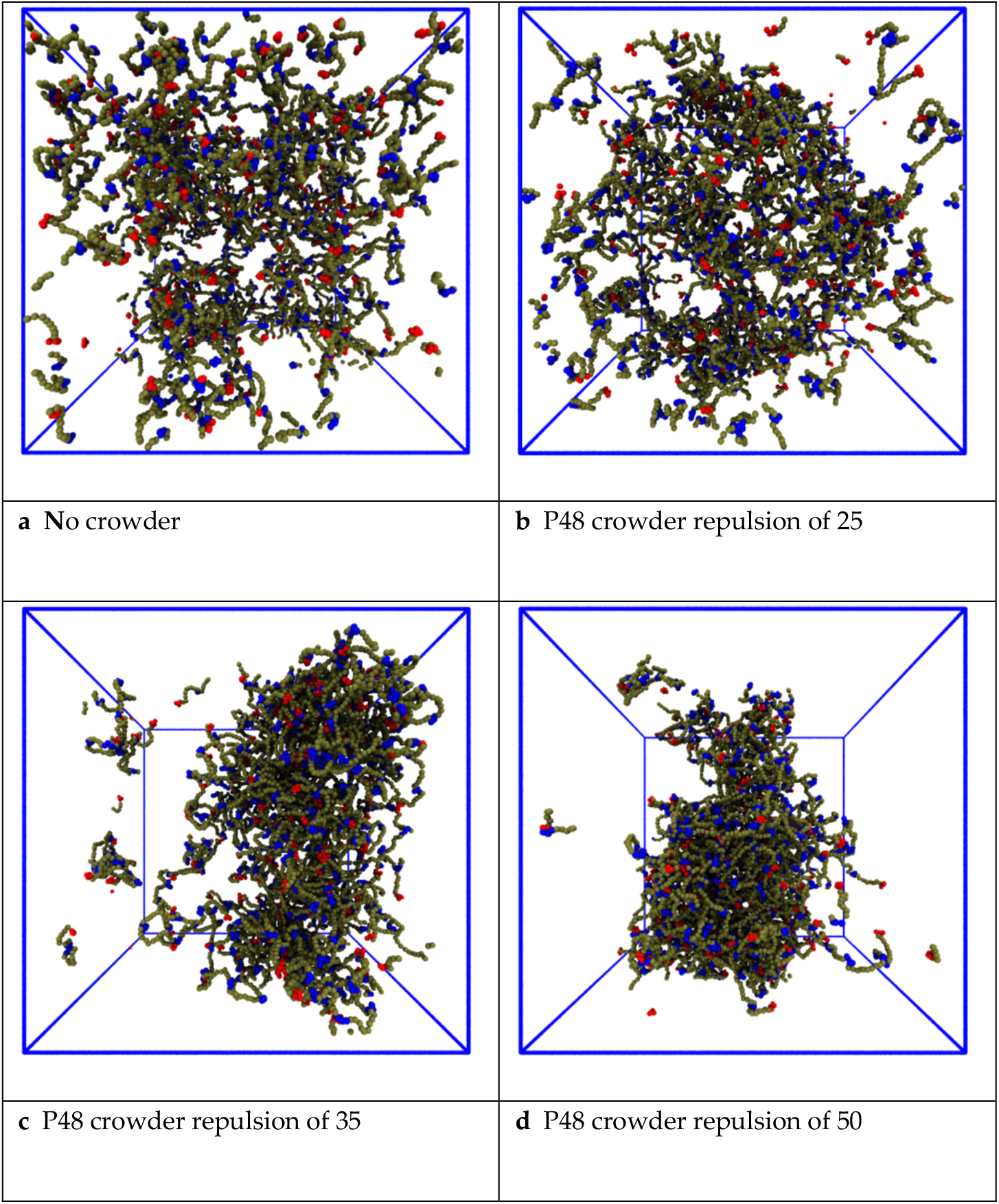
Effect of reducing the repulsive conservative force between the P48 crowder polymers and the IDPs. Each panel has the same number fraction of 6B10 IDPs with binding site affinity *ε* = 0.68. **(a)** The IDPs do not phase separate in the absence of crowder polymers. **(b)** The same repulsion between the crowders and the IDPs as their self-repulsion does not lead to phase separation. **(c)** A slightly greater repulsion enhances phase separation. **(d)** Stronger repulsion leads to complete phase separation.

## 4. Discussion

The cellular cytoplasm is a complex, crowded fluid, and understanding the effects of this environment on the phase separation of disordered proteins into biomolecular condensates is a challenging task. Most *in vitro* assays and computational models focus on a single protein, and their relevance to living cells is an open question.[39] A full statistical mechanical theory of the phase behaviour of a self-associating polymer appears extremely complex, although attempts of various kinds have been made.[65-67] In the presence of a polymeric crowding agent, it becomes (probably) impossibly complex. Yet experiments show that a crowded environment can influence the appearance of pathological fibrillar states within biomolecular condensates.[68-71] It has also been found that the material properties of BCs are important for their cellular functions,[34, 37] and disordered protein domains can exert steric pressure on cellular membranes.[72] Predicting the response of condensed phases of IDPs to the crowdedness of their environment is an important step in understanding their functions in health and possible role in treating diseases.[29, 30, 73]

Experiments on full-length FUS and its N-terminal low-complexity domain (FUS-LC) have shown that their dense phase is highly solvated.[42, 43] The flexible IDPs form transient attractive contacts along their length and retain large conformational fluctuations in the dense phase. LLPS (at least in the case of FUS-LC, which is almost uncharged and has fewer than 25% hydrophobic residues) is therefore unlike oil-water phase separation, and we do not expect mean field theories (including Flory Huggins theory) to describe their phase separation correctly.[74] Recent computational modelling suggests that conformational fluctuations are an important driver of the transition.[49, 50, 53, 75, 76]

In the present work, we explored the effects of adding a polymeric crowding agent on the phase separation of a model IDP using coarse-grained simulations. This provides a tractable model system to explore the range of possible responses of a BC to a crowded environment. The phase behaviour of the model proteins is controlled by their molecular weight, and the number, distribution, and affinity of their sticky sites. In the absence of a crowding agent, they phase separate into coexisting dilute and dense phases when the affinity of their sticky sites is sufficiently strong, form a space-filling network without phase separation for lower affinities, and remain dispersed when their attractive interactions are weak or absent.[45, 50, 56] The structural properties of the dense phase in similar coarse-grained models have been shown to be modulated by specific and non-specific molecular interactions.[77-80]

When a polymeric crowding agent is added to IDPs in solution, we observed the following effects: 1) if the IDP affinity is just below the value at which spontaneous phase separation occurs, the crowder shifts the transition so that the IDPs phase separate into a dense phase; 2) the observed dense phase has similar spatial structure to that of spontaneously-formed droplets of the same IDPs with higher affinity. The formation of the model biomolecular condensate is therefore coupled to the crowdedness of its environment, but its internal structure is largely decoupled, at least over the studied range. This may explain recent findings that the WNK1 kinase responds to osmotic stress in cells by condensing into a functional biomolecular condensate.[15] We note here that our simulations probe the situation in which crowding polymers only exert steric and entropic forces on the IDPs in a concentration range similar to the cellular cytoplasm. The densest system we simulated had a crowder concentration of 120 mM, which is comparable to the estimates of 100 – 450 mg/ml for cellular cytoplasm.[1, 2] This is a simpler case than the common crowding agent PEG, which strongly modifies the water activity in its environment.[3] Indeed, PEG has recently been observed not only to assist phase separation of the protein NPM1 with rRNA, but also to be concentrated inside the condensate.[81] This case could be explored by extending our approach to allow attractive interactions between the IDPs and the PEG polymers. Our results suggest that a cell may partition the cytoplasm into regions in which the phase boundary of a biomolecular condensate is tuned by the local concentration of external macromolecules (i.e., non-partitioning crowding agents) while its interior is (partially) decoupled from the cytoplasmic composition, thereby ensuring a stable internal fluid environment.

The spatial arrangement of the IDPs in the crowding-assisted dense phase is similar to that observed previously for the same IDPs with higher affinity that spontaneously phase separate. The structure of the model BC is therefore not strongly dependent on the strength of the attractive interactions between the constituent IDPs. We also observe that the model condensate still forms when the repulsion between the IDPs and the crowder is substantially lowered, and on reducing the crowder molecular weight keeping their volume fraction fixed.

We hypothesize that internal degrees of freedom of the IDPs, which are defined by their molecular sequence, are responsible for the robust structure of their dense phase. The punctate nature of their binding sites creates a spatial network of weak junctions connected by fluctuating spring-like lengths of polymer that resist deformation when the osmotic pressure due to the surrounding fluid increases. This may be compared to a single-component lipid bilayer vesicle, in which the membrane is stabilised by the strong hydrophobic repulsion of the lipid tails from the aqueous solvent. Adding cholesterol modifies the internal degrees of freedom of the lipids (chain ordering) and gives rise to a new phase – the liquid ordered phase. We speculate that as for the Lo phase, biomolecular condensates rely on the modification of their internal degrees of freedom to create robust microenvironments even in the presence of changing external concentrations of crowding agents.

Computational modelling provides a relatively inexpensive tool for exploring simplified representations of an experimental system, but suffers from its own complexity issues — namely that the number of parameters increases rapidly with increasing numbers of molecular species. Our exploration of LLPS in the presence of a crowding agent illustrates a powerful workflow for studying models with high-dimensional parameter spaces: 1) simultaneously generate many examples of a system at different points in its parameter space, and 2) rapidly compare different parts of the parameter space by viewing the data in a large grid of snapshots taken from different parameter space points, and identify the interesting regions. Tailoring this workflow for different systems is a promising step to overcoming the complexity gap between biological systems and simplified computational models. Further progress will likely involve using AI-enabled pattern recognition of the graphical arrays to identify the most interesting regions and direct costly compute resources towards the most efficient pathway of exploration.

## 5. Conclusions

Biomolecular condensates are widely viewed as providing a localised environment for cells to spatiotemporally segregate their biochemistry despite not being surrounded by a phospholipid membrane. The exchange of proteins between the dense and dilute phases and the crowding effect of other macromolecular species in the cytoplasm would appear to undermine their function. Our coarse-grained molecular simulations predict that a repulsive macromolecular crowding agent is able to drive a dilute solution of IDPs across their phase boundary into a dense phase even when their attractive self-interactions are too weak to drive phase separation alone. But, crucially, the spatial arrangement of proteins within the resulting BC is little changed from that of spontaneously-phase separating IDPs. These results also hold when molecular weight of the crowding polymer is reduced (by a factor of 2) providing their volume fraction is maintained, and when the crowder-IDP repulsion is reduced significantly. The dense phase structure is therefore predicted to be insensitive to the precise composition of its crowded environment. We propose that this arises because the pattern of attractive domains connected by entropic spring-like linkers along the IDPs creates a fluid, three-dimensional, network structure that resists deformation by the crowding polymers. Our results suggest that a cell may use the local cytoplasmic concentration of macromolecules to tune the formation of BCs, and that they may in turn sense, and respond to, changes in their environment.

When the self-affinity of the IDPs is removed, a sufficiently high concentration of the crowder drives a Flory-Huggins like demixing of the IDPs and crowder polymers, which visually resembles the model BC. However, the dense phase has no connected network structure, because the IDPs are unable to form transiently-stable junctions. When the IDP self-attraction is turned back on, the model BC reforms with the same internal structure as before (see Supplementary movies SM6 and SM7). This suggests that a functional BC may have a different spatial molecular arrangement than the same proteins compressed into a small volume solely by steric crowding.

Our prediction of two visually-similar dense phases, only one of which possesses a robust spatially-organised structure, implies that the observation of a demixing transition in computational studies of IDP phase separation is insufficient to conclude that their dense phase is a good model of a biomolecular condensate. The crucial question, we believe, is whether the spatial structure we observe, which is not reported in the literature by other groups, is found in experiments on biological condensates. Experimental tests of this prediction by positionally mutating key residues in the IDP FUS, and exploring the consequences for its phase behaviour in the presence of a crowder are under way.

## Author Contributions

Conceptualization, J.C.S., D.B.T., J.H.I., and A.D.B.; methodology, J.C.S., D.B.T.; software, J.C.S., D.B.T.; validation, J.C.S., D.B.T.; formal analysis, J.C.S. and D.B.T.; resources, A.D.B; writing—original draft preparation, J.C.S.; writing—review and editing, J.C.S, D.B.T., J. H. I., and A.D.B; visualization, J.C.S and D.B.T; supervision D.B.T, A.D.B; project administration, D.B.T and A.D.B; funding acquisition, A.D.B. All authors have read and agreed to the published version of the manuscript.

## Data Availability Statement

Simulation datasets are available on reasonable request to the corresponding author.

## Acknowledgments

The DPD source code used in this work is available on GitHub (https://github.com/Osprey-DPD/osprey-dpd)[62]. The authors acknowledge the use of the IRIDIS High Performance Computing Facility, and associated support services at the University of Southampton, in the completion of this work. The computational aspect of this research was supported by the EPSRC-funded POETS project (EP/N031768/1). Snapshots and movies of the simulations were produced using the open-source VMD software (http://www.ks.uiuc.edu/Research/vmd/)[82] and ImageJ (https://imagej.nih.gov/ij/index.html) [83].

