## Supplementary figures and images for "Macromolecular crowding is surprisingly unable to deform the structure of a model biomolecular condensate"

### Supplemental Figure 1

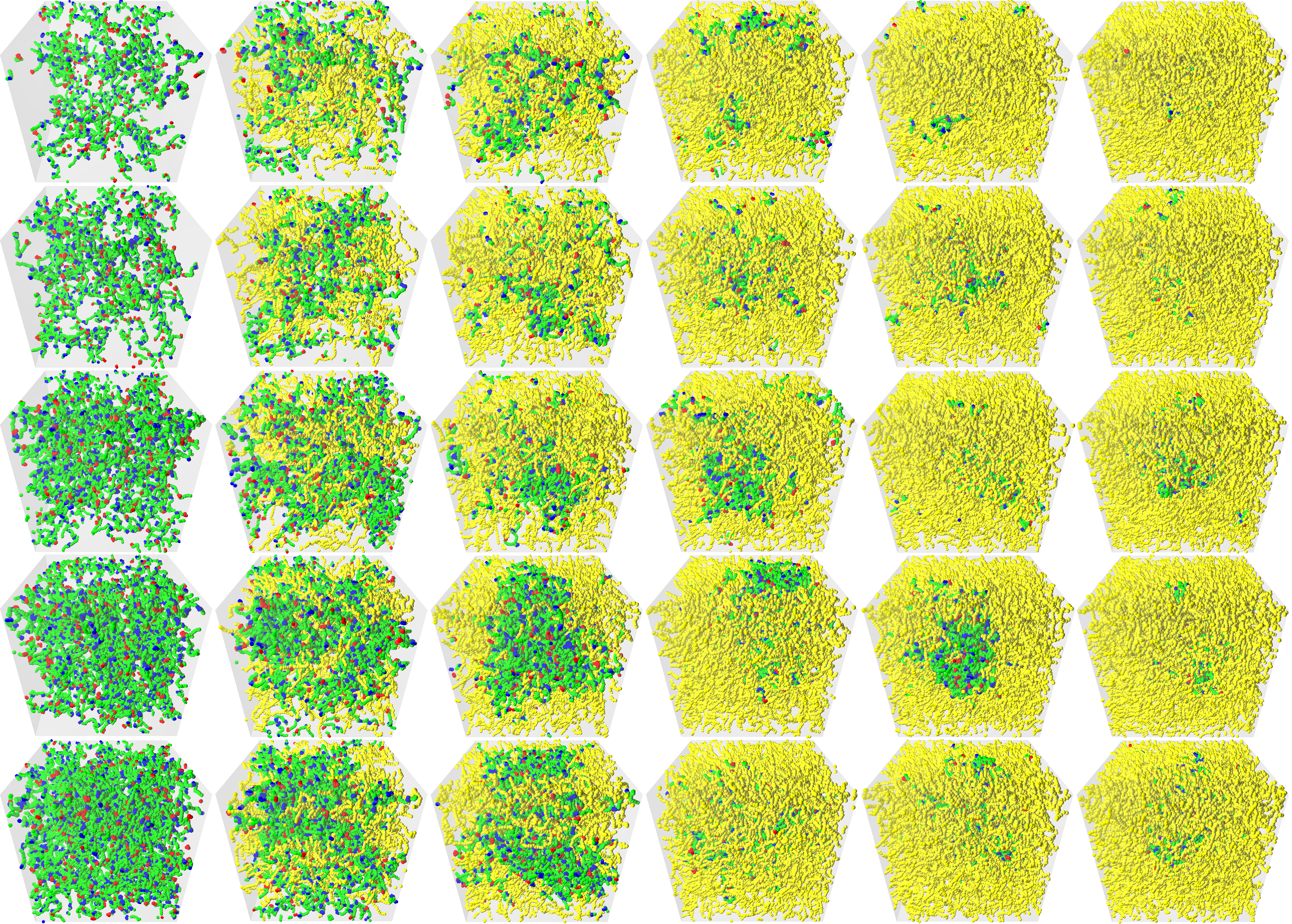

### Supplemental Figure 2

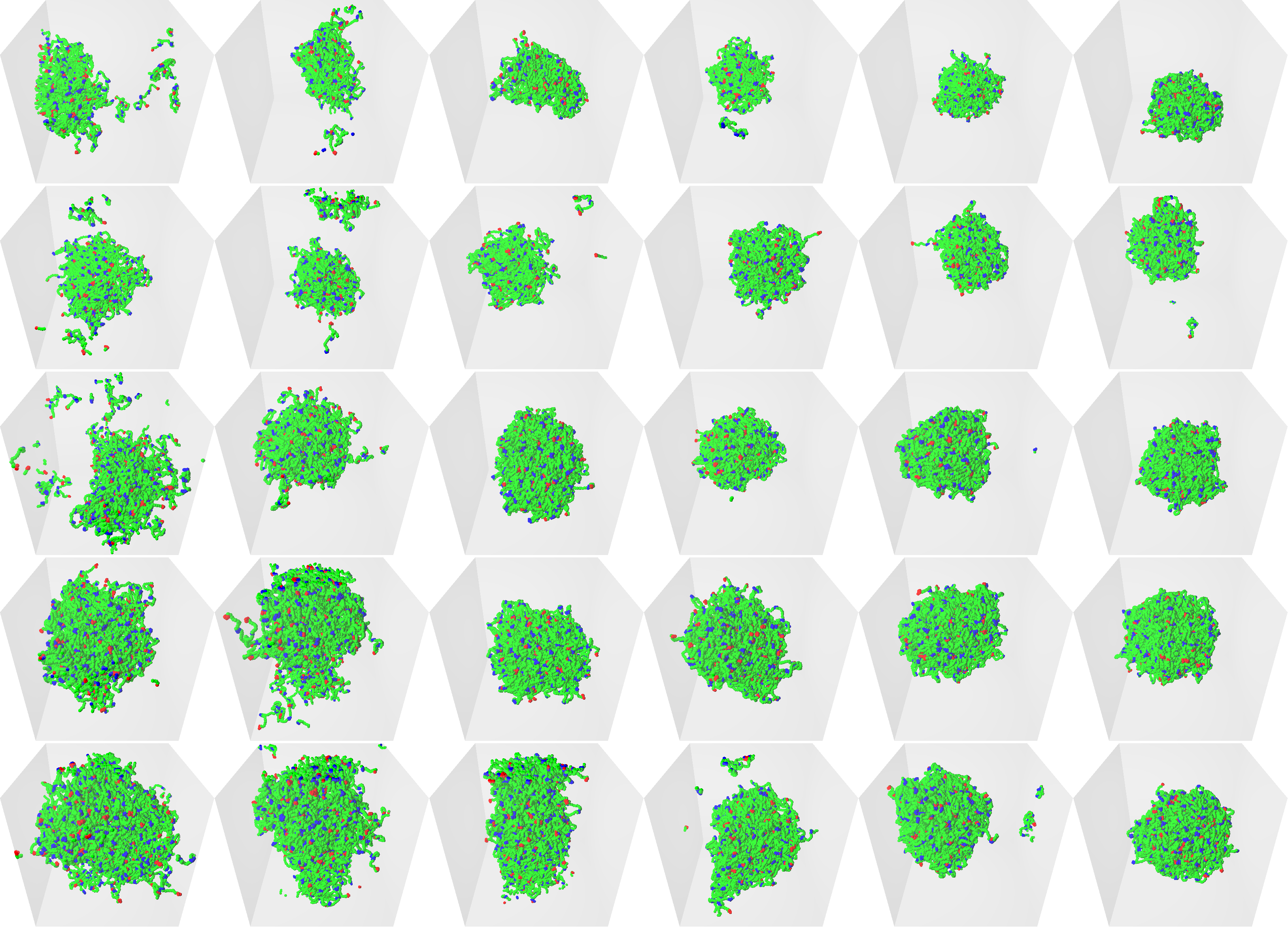

### Supplemental Figure 3

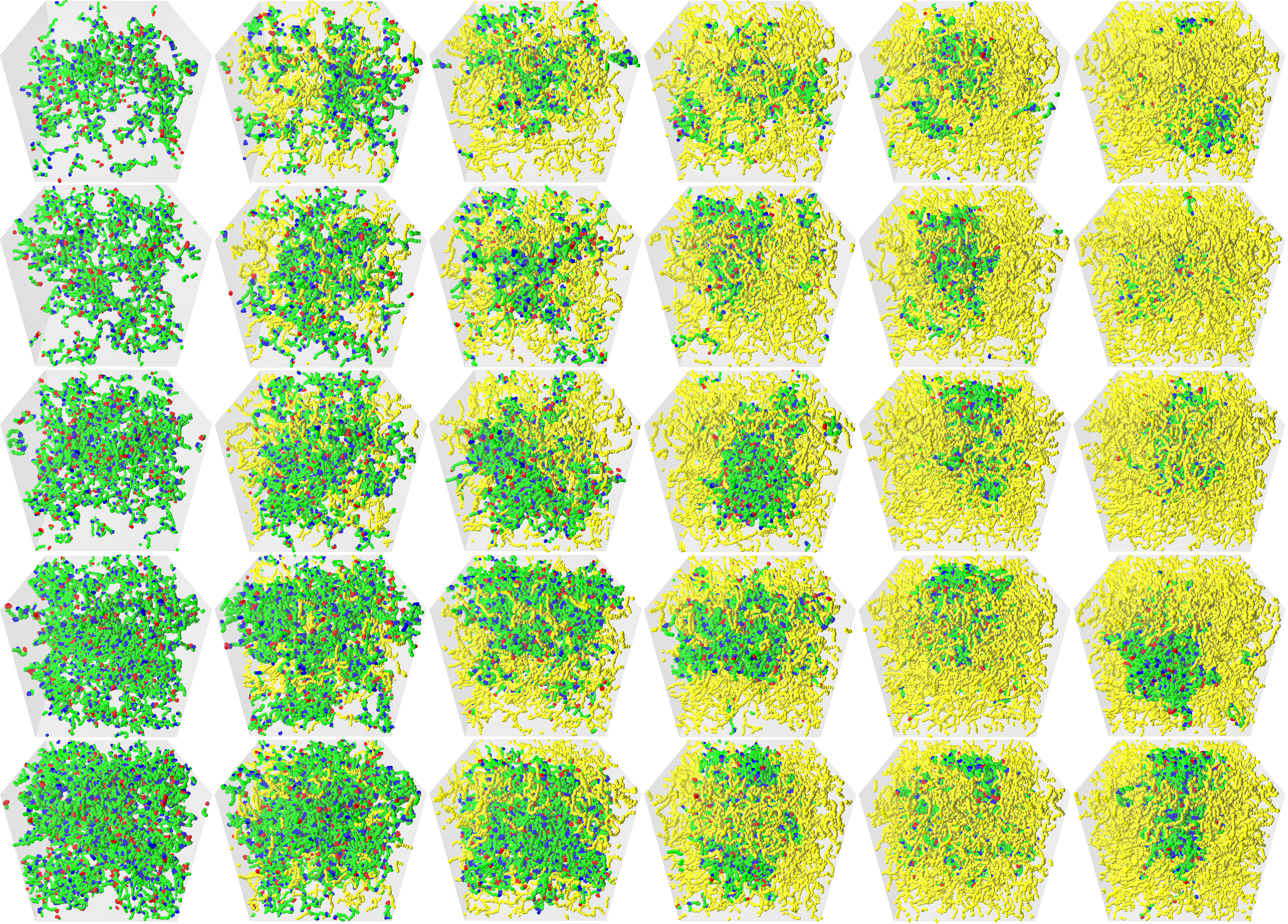
